# Oligonucleotide Length Determines Intracellular Stability of DNA-Wrapped Carbon Nanotubes

**DOI:** 10.1101/642413

**Authors:** Mitchell Gravely, Mohammad Moein Safaee, Daniel Roxbury

## Abstract

Non-covalent hybrids of single-stranded DNA and single-walled carbon nanotubes (SWCNTs) have demonstrated applications in biomedical imaging and sensing due to their enhanced biocompatibility and photostable, environmentally-responsive near-infrared (NIR) fluorescence. The fundamental properties of such DNA-SWCNTs have been studied to determine the correlative relationships between oligonucleotide sequence and length, SWCNT species, and the physical attributes of the resultant hybrids. However, intracellular environments introduce harsh conditions that can change the physical identities of the hybrid nanomaterials, thus altering their intrinsic optical properties. Here, through visible and NIR fluorescence imaging in addition to confocal Raman microscopy, we show that the oligonucleotide length determines the relative uptake, intracellular optical stability, and expulsion of DNA-SWCNTs in mammalian cells. While the absolute NIR fluorescence intensity of DNA-SWCNTs in murine macrophages increases with increasing oligonucleotide length (from 12 to 60 nucleotides), we found that shorter oligonucleotide DNA-SWCNTs undergo a greater magnitude of spectral shift and are more rapidly internalized and expelled from the cell after 24 hours. Furthermore, by labeling the DNA with a fluorophore that dequenches upon removal from the SWCNT surface, we found that shorter oligonucleotide strands are displaced from the SWCNT within the cell, altering the physical identity and changing the fate of the internalized nanomaterial. These findings provide fundamental understanding of the interactions between SWCNTs and live cells which can be applied towards development of robustly engineered carbon nanotube sensors while mitigating associated nanotoxicity.

**TOC Graphic.**
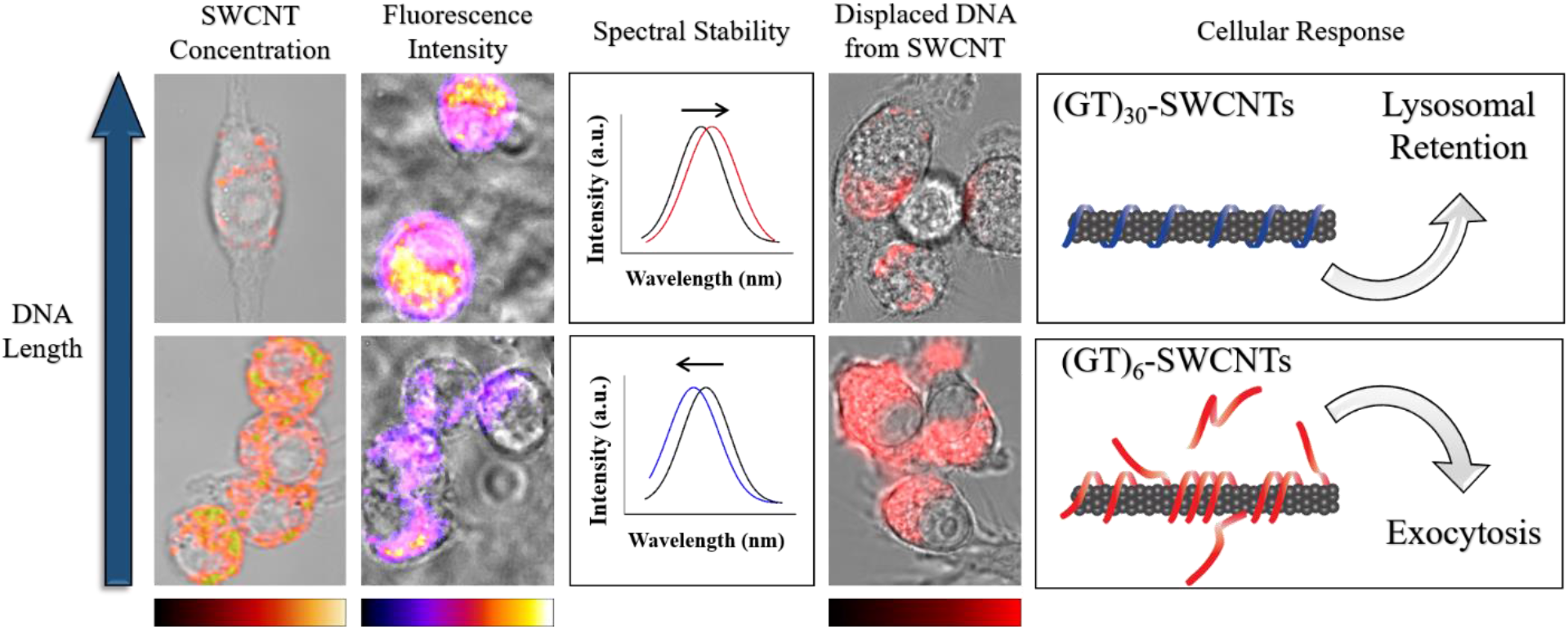

## Introduction

Single-walled carbon nanotubes (SWCNTs) have attracted substantial attention in the nanotechnology field due to their unique set of electrical [1], physical [2], and optical properties [3]. Their electronic band gap energies are dependent on their chiral identity, denoted by integers (*n,m)*, and vary based on diameter and rollup angle [4], resulting in semiconducting species which exhibit band gap photoluminescence [3]. Although highly hydrophobic in their raw as-produced form, non-covalent functionalization of SWCNTs using surfactants [5, 6] or amphiphilic biomolecules [7–9] has been shown to effectively disperse SWCNTs into aqueous solutions while preserving their intrinsic optical properties. Single-stranded DNA can non-covalently functionalize SWCNTs via π-stacking of hydrophobic bases onto the SWCNT sidewall, while the hydrophilic phosphate backbone allows for significantly enhanced aqueous solubility [10]. These DNA-SWCNT hybrids have shown promise as biological imaging [11] and sensing probes [12] due to their near-infrared (NIR) photoluminescence which is tunable, photostable, and sensitive to their local environment [13–16].

Hybrids of DNA and SWCNTs are preferred over other non-covalent approaches due to their enhanced biocompatibility [17], ability to sort single (*n*,*m*)-chiralities from parent mixtures [18, 19], and the potential for sensing imparted by the inherent diversity of oligonucleotide sequence. Specific sequence formulations of DNA-SWCNTs have been recently used to detect miRNA *in vivo* [20] in addition to reporting lipid concentrations in live cells [21] and animals [22], however fundamental questions relating the identity of these sensors after prolonged exposure within the biological environment remain largely unexplored. The potential instability of such DNA-SWCNT sensors has direct implications on their ability to perform a designated task, yet the indirect consequence is a nanomaterial with altered properties from its original state, thus leading to concerns about biological impact and toxicity. While many types of DNA-SWCNTs have been studied extensively *in situ* both computationally [23–25] and experimentally [26–31], their direct translation to more complex biological systems cannot be assumed.

Macrophages are the immune system’s first line of defense, whether as a primary response to a wound or to engulf foreign substances that enter the bloodstream. Various studies have shown that macrophages internalize DNA-SWCNTs via endocytosis and phagocytosis through the endolysosomal pathway, eventually leading to localization within the lysosomes [21, 32, 33] and accumulation in the liver macrophages of mice *in vivo* [22, 34]. Once entrapped within the lysosomes, SWCNTs can remain for days where they experience biologically low pH and exposure to more than 60 hydrolases meant for catabolic degradation [35]. In these conditions, surface modifications can play a large role on a nanoparticle’s ultimate fate, whether degradation, exocytosis, or lysosomal escape [36]. Given their extremely high surface area to volume ratio, small changes in surface functionalization of SWCNTs can make a major impact on their functionality and stability in such environments.

While oligonucleotide length determines the intrinsic stability of the resultant hybrid with a SWCNT in water [28], little is known how this length of DNA can affect the stability of SWCNTs in complex intracellular environments. Herein, we present an investigation of the physical and optical stabilities of (GT)_n_-SWCNTs, where n is the number of sequence repeats, upon internalization into murine macrophages. Near-infrared hyperspectral microscopy in live cells revealed strong correlations between oligonucleotide length and the NIR fluorescence intensity and spectral stability of the examined SWCNTs. All DNA-SWCNT combinations displayed emission shifts to lower energies (i.e. red-shifts) upon interacting with the cells, however smaller diameter (GT)_6_-SWCNTs exhibited significant blue-shifts over the course of 24 hours, indicating molecular adsorption and/or DNA displacement. We quantified SWCNT concentrations in cells using confocal Raman microscopy and revealed significant differences in both internalization and expulsion of (GT)_6_- and (GT)_30_-SWCNTs over 24 hours. Finally, we used fluorophore labeled DNA to probe the condition of the SWCNT hybrids as they were processed through the endolysosomal pathway.

## Results and Discussion

To study the effects of single-stranded DNA length on the intracellular optical properties of DNA-SWCNTs, we first non-covalently functionalized HiPco SWCNTs with one of five different (GT)_n_ oligonucleotides, where *n* = 6, 9, 12, 15, or 30 repeats (Fig. S1). Murine macrophages (RAW 264.7 cell line) were pulsed for 30 minutes with 1 mg/L of each (GT)_n_-SWCNT sample under standard cell culture conditions and the majority of cells exhibited substantial NIR broadband fluorescence (ca. 900-1600 nm) when excited by a 730 nm laser (Fig. 1a). In agreement with previous studies [15, 21, 37], NIR fluorescence movies confirmed the internalization of the SWCNTs into endosomal vesicles, which were actively translocated around the cell (Movie S1). The NIR fluorescence images were acquired 0, 6, or 24 hours after an initial pulse to assess the DNA length and temporal dependencies on intracellular fluorescence intensity (Fig. 1b). In general, the observed NIR fluorescence intensities visibly increased with increasing oligonucleotide length, but decreased in time after initial loading into the cells. Histograms constructed from pixel intensity values of the 0- and 24-hour images confirmed that the temporal decreases in intensities were similar amongst all sequences (Fig. 1c). Interestingly, the initial intensity distributions were much broader in longer oligonucleotide sequences, suggesting more heterogeneity in the optical response of these SWCNTs. To quantify the images, the average fluorescence intensities were extracted using a global thresholding analysis to examine the NIR fluorescence from only SWCNTs contained within the cells. We observed significant increases in observed NIR fluorescence intensities as a function of DNA length (Fig. 1d). While this finding is in agreement with relative quantum yield values determined from solution measurements (Fig. S1d), we propose that these results are also affected by (1) variations in DNA-SWCNT interaction with and internalization into cells as a function of DNA length or (2) variations in the optical stability of the (GT)_n_-SWCNT hybrids after interacting and/or internalizing into the cells. Throughout the manuscript, we will carefully examine these hypotheses.

**Figure 1:**
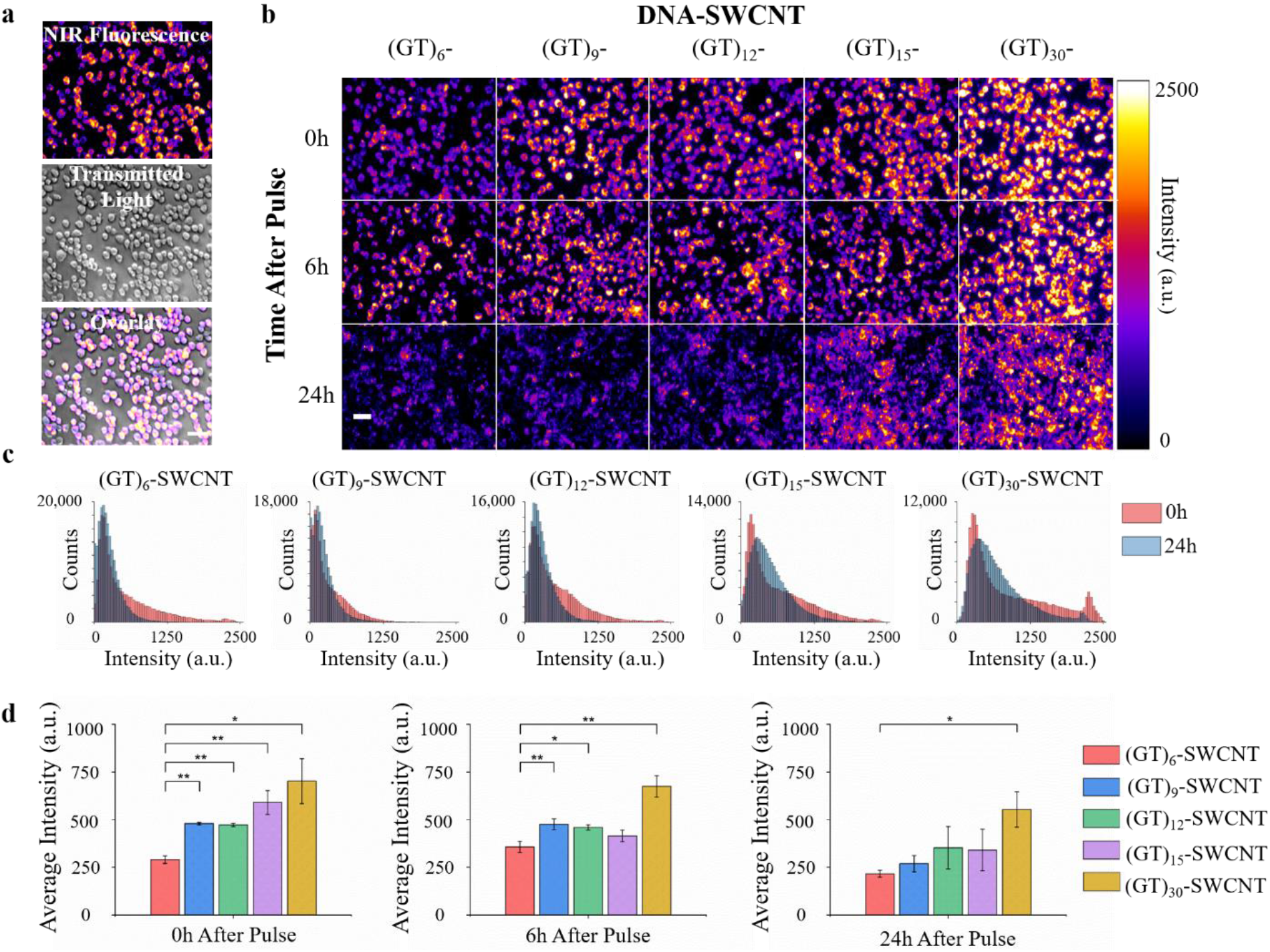
Length-dependent intracellular fluorescence of DNA-SWCNTs. **(a)** NIR fluorescence image of live macrophages pulsed with (GT)_15_-SWCNTs, along with respective transmitted light image and merged NIR/transmitted light image. **(b)** NIR fluorescence images of macrophages after 30-minute pulse of (GT)_n_-SWCNTs, imaged over the course of 24 hours. Scale bar = 20μm. **(c)** Histograms corresponding to the 0- and 24-hour (GT)_n_-SWCNT images in (b), and **(d)** average intracellular fluorescence intensities for all examined DNA sequences 0-, 6-, or 24-hours after (GT)_n_-SWCNT pulse. Experiments were performed in triplicate and are represented as mean ± s.d. (*, p < 0.05, **, p < 0.01, according to two-tailed two-sample t-test).

We employed NIR hyperspectral fluorescence microscopy to assess the chirality-resolved intracellular stability of the (GT)_n_-SWCNTs [15]. Using a 730 nm excitation laser, we were able to resolve four distinct bands in the NIR region corresponding to the emission spectra of the four brightest SWCNT chiralities, (10,2), (9,4), (8,6), and (8,7) (Fig. S1a,b) [15]. Hyperspectral images were acquired immediately following a 30-minute pulse of each (GT)_n_-SWCNT and after an additional 24 hours of incubation in SWCNT-free cell media (Fig. 2a). Upon internalization, we observed two common characteristics of all fluorescence spectra: (1) an initial red-shift (i.e. increase in wavelength) of every chirality compared to the spectra aquired in cell culture media (Fig. S1b) and (2) increased intensities of longer wavelength chiralities relative to shorter. To explain the first finding, a red-shift in SWCNT emission spectra can be caused by charged species that interact with the phosphate backbone of DNA and induce a conformational change, ultimately modulating the dielectric environment of the SWCNT and thus shifting SWCNT emission to longer wavelengths [38]. Surface proteins present on cell membranes with high charge densities have been shown to promote this red-shift upon first contact with DNA-SWCNTs before endocytosis [37]. Additionally, the exposure of DNA-SWCNTs to serum-containing cell culture media can produce aggregation and spectral modulation by protein-DNA electrostatic interactions [39]. We attribute the initial red-shift observed to a combination of these factors directly following a pulse of (GT)_n_-SWCNTs, in which the macrophages contained both membrane bound particles that had not yet been internalized as well as newly-formed endosomal vesicles that essentially forced DNA-SWCNTs to form small aggregate complexes with other phagocytosed proteins and cargo. Regarding the second finding, changes in the ratiometric intensities between shorter and longer emission wavelength SWCNTs have been described by inter-nanotube exciton energy transfer (INEET) [40], a phenomenon that behaves similarly to Förster resonance energy transfer and could be the result of closely packed DNA-SWCNTs contained within lysosomes.

**Figure 2:**
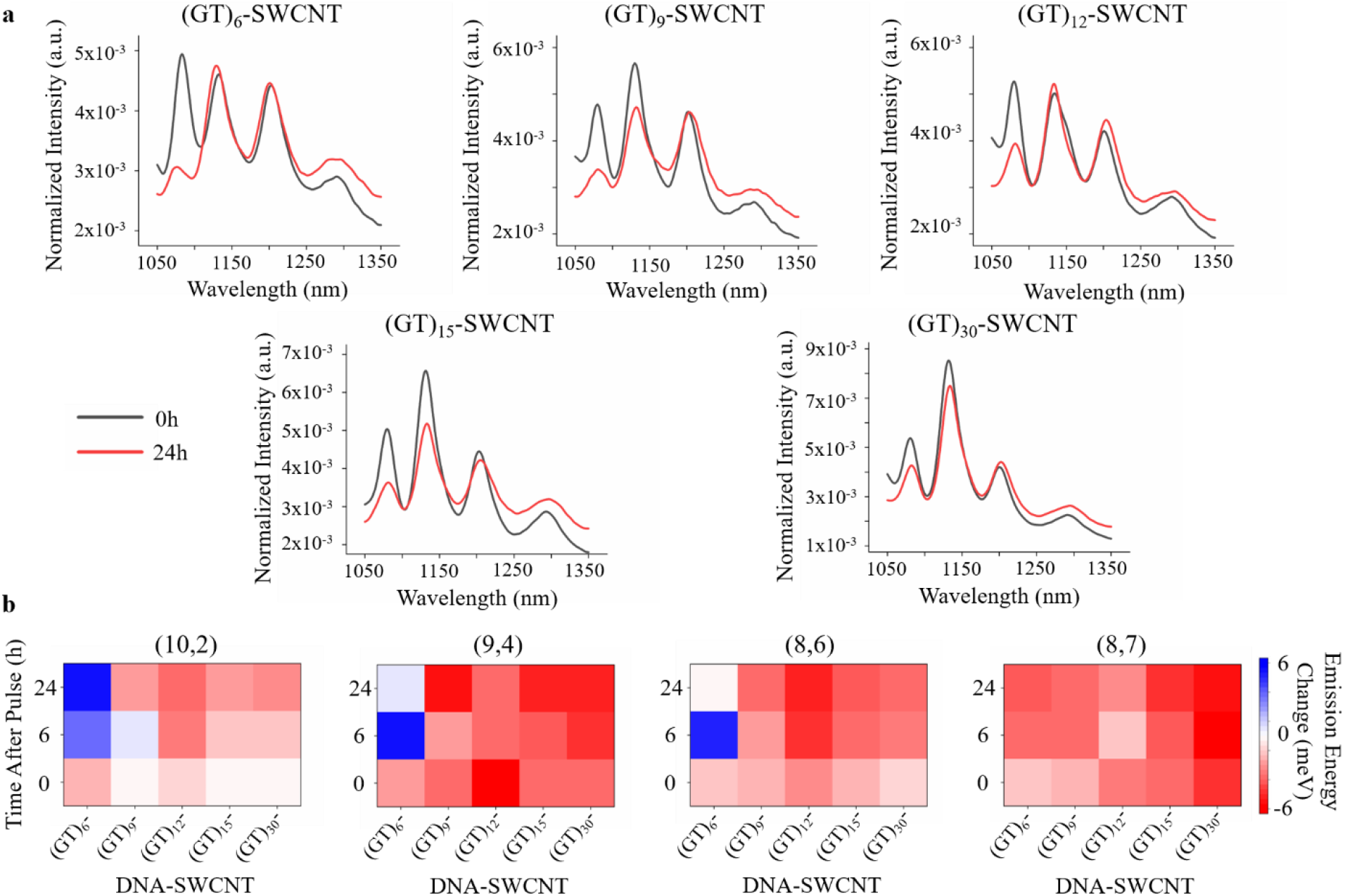
DNA length- and SWCNT chirality-dependent intracellular NIR fluorescence stabilities. **(a)** Intracellular fluorescence spectra 0- and 24-hours after internalization for each (GT)_n_-SWCNT. The intensity of each spectra was normalized to the area under the curve. **(b)** Heat maps representing intracellular change in SWCNT emission energy compared to controls in cell culture media, delineated by chirality, as a function of DNA-sequence and time. All experiments were performed in triplicate.

To further compare the NIR fluorescence stabilities across SWCNT chiralities, we converted the emission center wavelengths to energies (meV) and computed the change in emission energy relative to solution controls acquired in cell culture media. Of all the examined oligonucleotide lengths, only the (GT)_6_-SWCNTs exhibited significant increases in emission energies in multiple chiralities over 24 hours of intracellular processing (Fig. 2b). Previous studies have demonstrated that this increase in emission energy could arise from endosomal lipids binding to the exposed SWCNT surface [21, 22], however each chirality temporally displayed variable emission shifts within the same population of cells, indicating the reported shifts could be due to more than one factor. The other DNA-SWCNTs commonly displayed a moderate loss of energy over the same period of time despite identical intracellular conditions. We believe this indicates that a longer oligonucleotide wrapping protects the SWCNT surface from competitive molecular adsorption.

To assess spectral shifts in SWCNT emission at the single-cell level, we created hyperspectral maps of the shortest and longest oligonucleotide DNA-SWCNTs (i.e. (GT)_6_ and (GT)_30_, respectively). By fitting each SWCNT-containing pixel of a hyperspectral image to a Gaussian curve [15, 21], we were able to overlay transmitted light images with center emission energy maps for the (9,4)-SWCNT, i.e. the most abundant and brightest SWCNT under 730 nm excitation in HiPco, and construct histograms for each image to depict the intracellular change in SWCNT emission energy change through time (Fig. 3a,b). Immediately following a 30-minute pulse (“0 hours”), the average emission energies of (GT)_6_-SWCNTs and (GT)_30_-SWCNTs were statistically identical. Additionally, by fitting the pixel histograms to a Gaussian distribution, the heterogeneity in the populations could be assessed by examining the full width at half maximum (FWHM). In doing so, we uncovered that the FWHM of (GT)_6_-SWCNTs immediately after internalization was more than double that of (GT)_30_-SWCNTs. While the emission energy of (GT)_30_-SWCNTs showed little change in time, (GT)_6_-SWCNTs displayed an 8 meV increase in emission energy and ~50% decrease in FWHM after 6 hours. We believe these DNA-length dependent NIR fluorescence modulations are the result of variations in the relative abundance of oligonucleotide strand ends surrounding each SWCNT. For a given weight of DNA in a DNA-SWCNT hybrid, (GT)_6_-SWCNTs have 5 times the number of oligonucleotide strand ends than (GT)_30_-SWCNTs. We propose that these strand ends can act as initiation sites for amphiphilic biomolecules to interact with and adsorb onto the exposed nanotube surfaces, leading to higher overall surface coverages (reduced water densities) and thus greater emission energies [21, 41]. Consequently, the wide FWHM initially displayed by (GT)_6_-SWCNTs was likely caused by individual SWCNTs responding to the varying local environments through progressing stages of the phagocytic pathway, while the reduced FWHM and blue-shift after 6 hours can be attributed to molecular adsorption by lysosomal molecules and rearrangement or displacement of the oligonucleotide wrapping on the majority of SWCNTs. Interestingly, after 24 hours the emission energy slightly decreased closer to its initial value while the FWHM increased towards its initial value, revealing that the hybridized (GT)_6_-SWCNTs observed at 6 hours were ultimately unstable.

**Figure 3:**
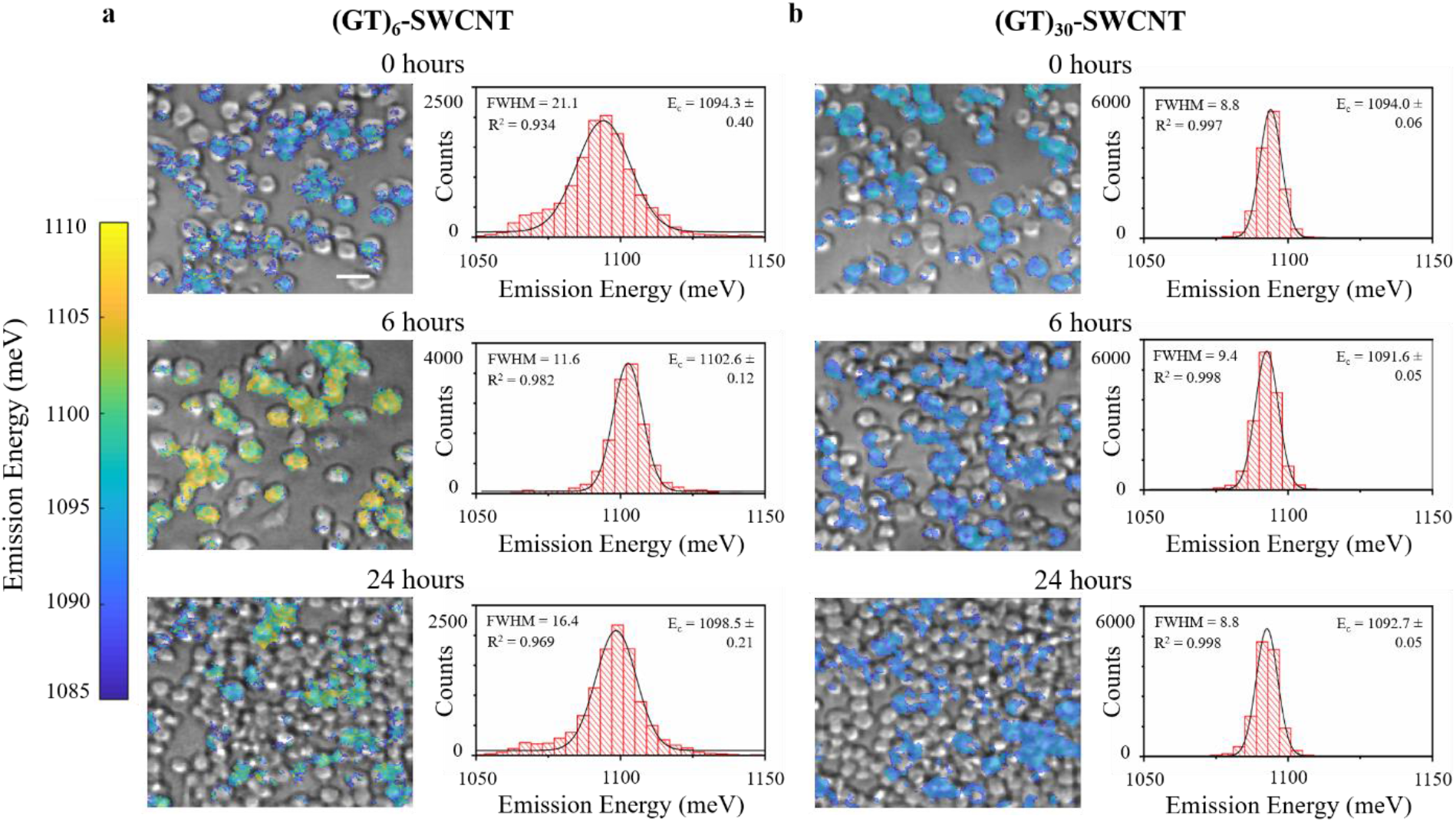
Hyperspectral maps of DNA-(9,4)-SWCNTs in live macrophages. Overlay of transmitted light and hyperspectral images of RAW 264.7 pulsed for 30-minutes with **(a)** (GT)_6_- or **(b)** (GT)_30_-SWCNTs. Color scale maps to the fitted center emission energy of (9,4)-SWCNTs and histograms represent center emission energies of all SWCNT-containing pixels in each respective image. Bin size = 4 meV. Gaussian functions were fitted to binned data and overlaid with respective R^2^, FWHM, and E_c_. Scale bar = 20 μm.

While the observed fluorescence intensities of SWCNTs can be modulated by both concentration and local-environment [16, 41, 42], certain Raman signatures of SWCNTs depend only on concentration [43–46]. Therefore, we assessed the localized intracellular concentrations of SWCNTs using confocal Raman microscopy. Small regions were scanned in 0.5 μm intervals to obtain Raman maps of macrophages pulsed with 1 mg/L (GT)_6_- or (GT)_30_-SWCNTs (Fig. 4a). The intensity of the G-band spectral feature, indicative of *sp*^2^ carbon [44, 45], was correlated to known SWCNT concentrations in the construction of a calibration curve in order to obtain a mass of SWCNTs per analyzed cell (Fig. S2). Although the local concentrations varied greatly within a single cell, on average the cells pulsed with (GT)_6_-SWCNTs had more than twice the initial intracellular SWCNT weight than those incubated with (GT)_30_-SWCNTs (Fig. 4b). The higher uptake of (GT)_6_-SWCNTs could be caused by their higher overall density of DNA per SWCNT compared to (GT)_30_-SWCNTs [26], increasing the probability of interactions between DNA and cellular membrane proteins, thus leading to more nanotubes per engulfing phagosome. After 24 hours of additional incubation in SWCNT-free cell media, the internal SWCNT concentration of cells dosed with (GT)_6_-SWCNTs dropped more than 75%, while those dosed with (GT)_30_-SWCNTs displayed statistically similar initial and final concentrations, indicating that the effects of intracellular processing on DNA-SWCNTs are determined by oligonucleotide length. We surmise that changes in the physical identity of internalized (GT)_6_-SWCNTs are inducing the macrophages to exocytose this sample more rapidly than the stable (GT)_30_-SWCNTs. We further propose that integrity of the DNA-SWCNT hybrids is the key factor in determining whether or not a cell chooses to exocytose the nanotubes.

**Figure 4:**
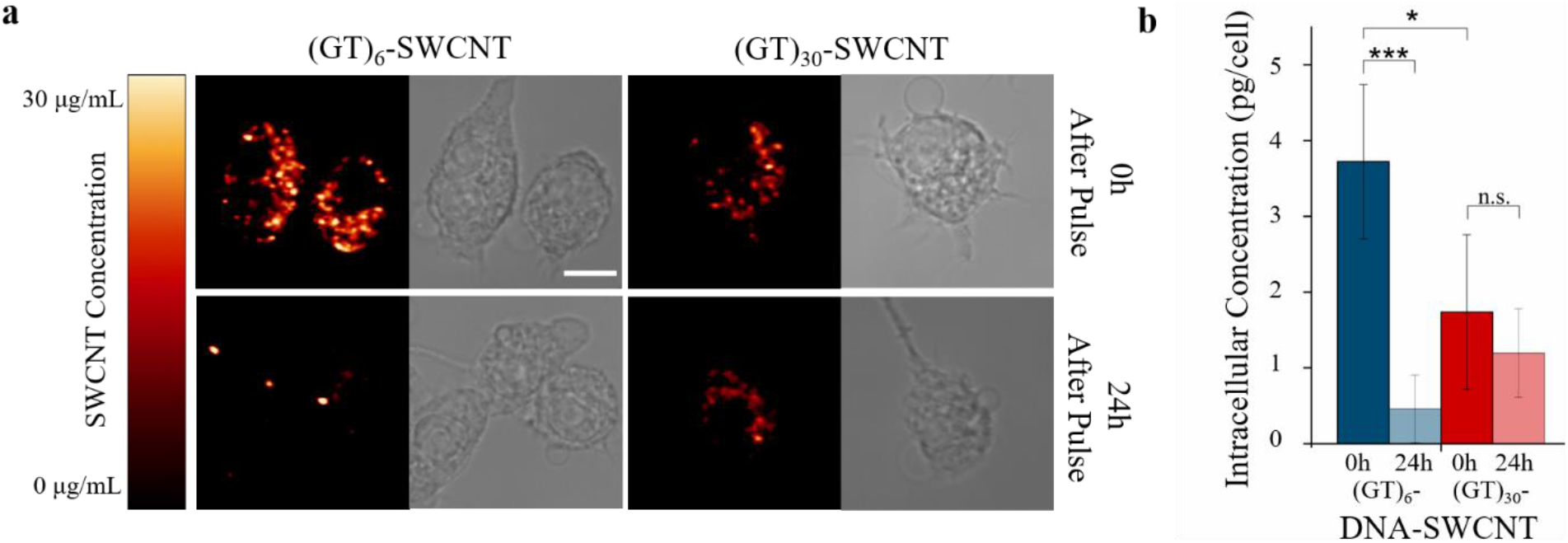
SWCNT concentration maps determined by confocal Raman microscopy. **(a)** Confocal Raman microscopy images showing G-band intensity and white light images of RAW 264.7 cells pulsed with (GT)_6_- or (GT)_30_-SWCNTs for 30 minutes. Color map represents SWCNT pixel concentration. Scale bar = 10 μm. **(b)** Average SWCNT concentration (n ≥ 4 cells) calculated from total pixel concentration within cellular ROIs. Error bars represent mean ± s.d. (*P < 0.05, ***P < 0.001, according to two-tailed two-sample t-test).

To further probe the integrity of the (GT)_n_-SWCNT hybrids while contained by macrophages, we devised an assay based on the ability of SWCNTs to quench conventional organic fluorophores [47]. We first constructed (GT)_6_- or (GT)_30_-SWCNTs with a Cy3 dye attached to the 5’ end of the DNA strand. The initially quenched fluorophore could be restored to a brightly fluorescent state via displacement from the SWCNT surface by a competing molecule (Fig. 5a). Note, even partial displacement of the DNA strand can accomplish this process, thus Cy3 dequenching kinetics of the two prepared hybrids are similar (Fig. 5b) despite unequal displacement kinetics [28]. When introduced to macrophage cells in the same 30-minute pulse method, we observed substantially different dequenching behavior between the two sequences (Fig. 5c). The Cy3-(GT)_6_-SWCNTs significantly dequenched inside of the cells, reaching a maximum intensity 4 hours after internalization (Fig. 5d) and decreasing to its initial intensity after 24 hours. In contrast, dequenching was not observed in the Cy3-(GT)_30_-SWCNTs at any point, resulting in statistically significant differences in the dequenching ability of the DNA-SWCNTs within the first six hours.

**Figure 5:**
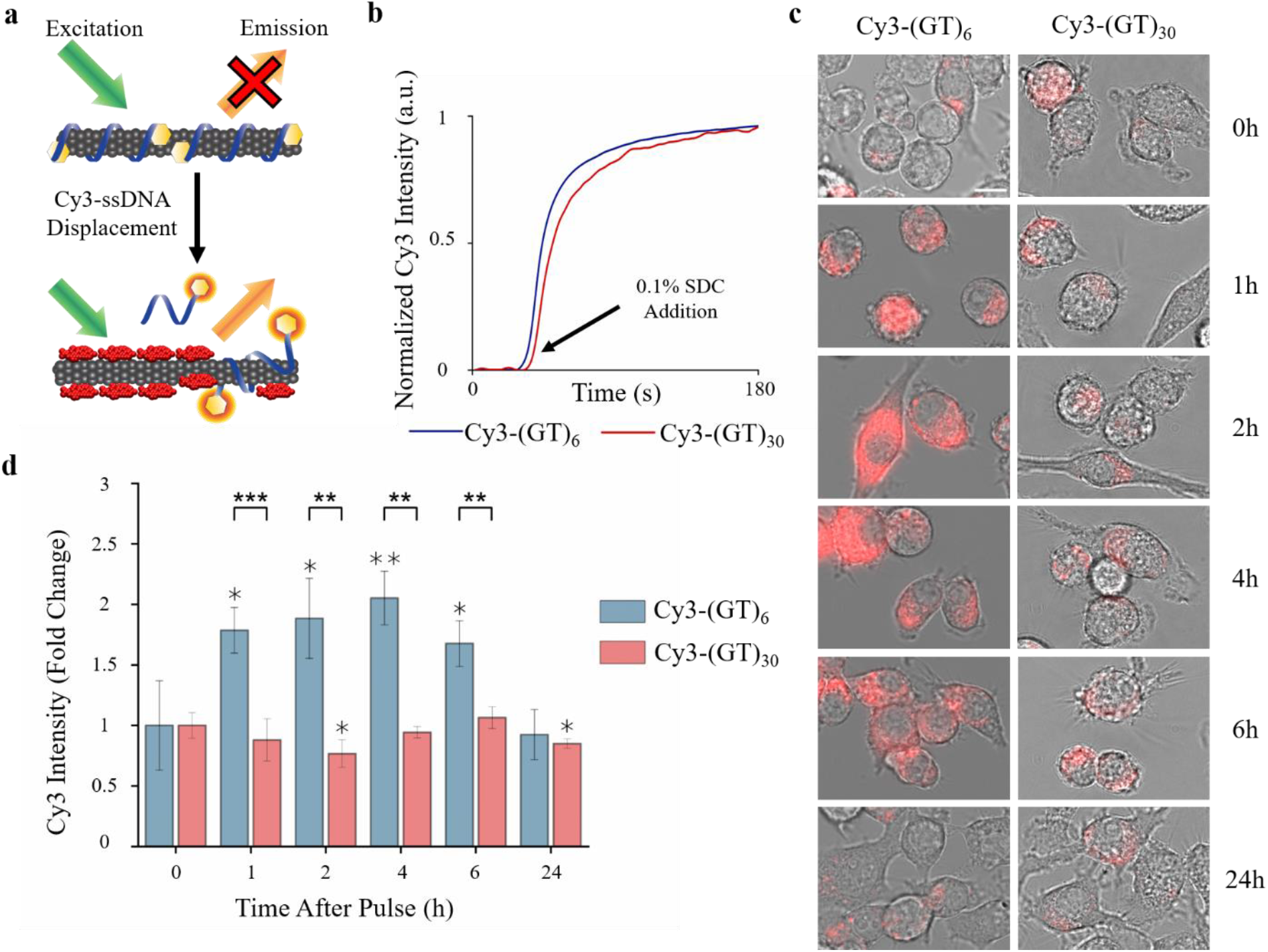
Intracellular stability of DNA-SWCNT hybrids. **(a)** Schematic of experimental design. Cy3-DNA is quenched when wrapping is intact on SWCNTs, but highly fluorescent once displaced. **(b)** Normalized intensity increase as a function of time after Cy3-DNA is displaced with SDC. **(c)** Overlaid Cy3-DNA and white light images of RAW 264.7 pulsed with Cy3-(GT)_6_ or Cy3-(GT)_30_-SWCNTs for 30 minutes. Scale bar = 10 μm. **(d)** Average fluorescence intensities (n ≥ 14 cells) normalized to 0-hour intensity. Error bars represent mean ± s.d. Five-pointed stars represent significance between Cy3-(GT)_6_ and Cy3-(GT)_30_ and six pointed stars represent significance versus initial intensities. (*P < 0.05, **P < 0.01, ***P < 0.001 according to two-tailed two-sample t-test).

## Conclusions

We propose that the intracellular processing and ultimate fate of (GT)_n_-SWCNTs are determined by the differential stabilities of the hybrid nanomaterials in the lysosomal environment, which correlate strongly to the length of a given DNA strand. The observed intracellular fluorescence intensities were shown to increase with increasing oligonucleotide length, while only the shortest DNA-SWCNTs (i.e. (GT)_6_) displayed instabilities in NIR fluorescence spectra in time. We have shown that (GT)_30_-SWCNTs are mostly retained within the cells over 24 hours with minimal exocytosis, while (GT)_6_-SWCNTs expelled more than 75% of the internalized cargo over the same time period despite nearly a two-fold higher amount of initial uptake. Furthermore, we demonstrated that (GT)_6_ is highly susceptible to displacement or degradation from the SWCNTs by utilizing a fluorophore dequenching assay, providing further support that the intracellular stability of the DNA-wrapping on SWCNTs depends strongly on oligonucleotide length. Therefore, we propose a schematic (Fig. 6) to describe the series of events that occur to (GT)_n_-SWCNTs upon internalization by macrophages and accentuate the stabilities of the resultant hybrids as well as the cellular responses due to these instabilities. These results provide important insights for DNA-SWCNT-based sensor design at the cellular level and highlight the sensitivity of nanoscale systems to surface modifications in biological environments.

**Figure 6:**
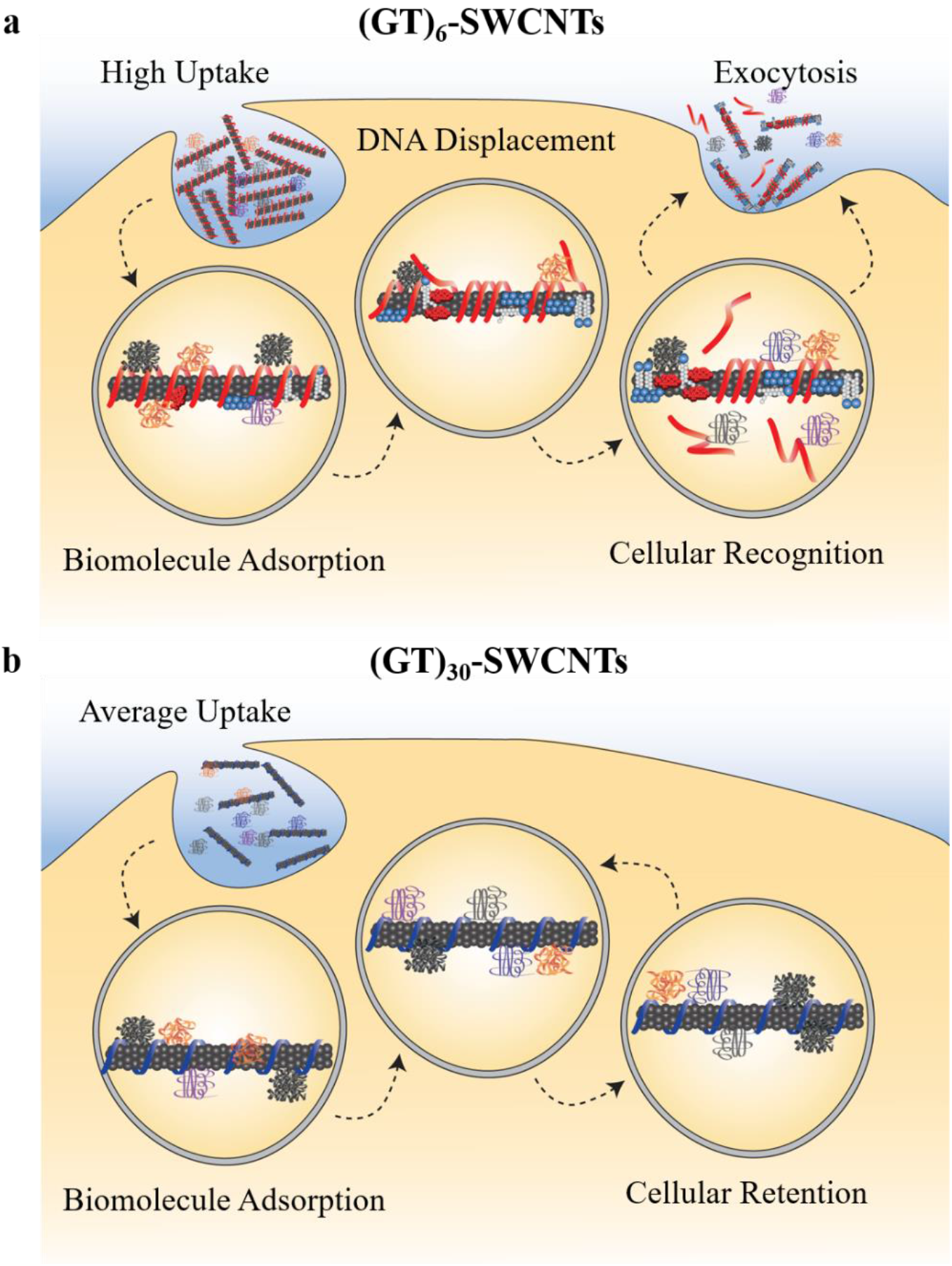
Schematic depicting the DNA length-dependent intracellular processing of DNA-SWCNTs. **(a)** Short-sequence (GT)_6_-SWCNTs are internalized in large amounts and localized to the lysosomes, where biomolecules can adsorb to the nanotube surface before eventually inducing DNA displacement and ultimately leading to exocytosis. **(b)** Long-sequence (GT)_30_-SWCNTs are also located to the lysosomes after internalization but do not experience molecular adsorption directly to the nanotube surface, preventing DNA displacement and reducing exocytosis.

## Materials and Methods

### DNA-SWCNT Sample Preparation

Raw single-walled carbon nanotubes produced by the HiPco process (Nanointegris) were used throughout this study. For each dispersion, 1 mg of raw nanotubes was added to 2 mg of (GT)_n_ (where *n* = 6, 9, 12, 15, or 30) oligonucleotide (Integrated DNA Technologies), suspended in 1 mL of 0.1M NaCl (Sigma-Aldrich), and ultrasonicated using a 1/8” tapered microtip for 30 min at 40% amplitude (Sonics Vibracell VCX-130; Sonics and Materials). The resultant suspensions were ultra-centrifuged (Sorvall Discovery M120 SE) for 30 min at 250,000 x*g* and the supernatant was collected. Concentrations were determined using a UV/vis/NIR spectrophotometer (Jasco, Tokyo, Japan) and the extinction coefficient of A_910_ = 0.02554 L mg^−1^ cm^−1^ [15].

### Cell Culture

RAW 264.7 TIB-71 cells (ATCC, Manassas, VA, USA) were cultured under standard incubation conditions at 37 °C and 5% CO_2_ in cell culture medium containing sterile filtered high-glucose DMEM with 10% heat-inactivated FBS, 2.5% HEPES, 1% L-glutamine, 1% penicillin/ streptomycin, and 1% amphotericin B (all acquired from Gibco). For all cell-related studies, cells were allowed to grow until 90% confluency and used up to the 20^th^ passage.

### Near-Infrared Fluorescence Microscopy of Live Cells

A near-infrared hyperspectral fluorescence microscope, similar to a previously described system [15], was used to obtain fluorescence images and hyperspectral data within live cells. In short, a continuous 730 nm diode laser with 1.5 W output power was injected into a multimode fiber to produce an excitation source, which was reflected on the sample stage of an Olympus IX-73 inverted microscope equipped with a 20X LCPlan N, 20x/0.45 IR objective (Olympus, USA) and a stage incubator (Okolab) to maintain 37 °C and 5% CO_2_ during imaging. Emission was passed through a volume Bragg Grating and collected with a 2D InGaAs array detector (Photon Etc.) to generate spectral image stacks. For live cell experiments, cells were seeded into tissue culture treated 96-well plates (Fisher Scientific) at a final concentration of 25,000 cells/ well and allowed to culture overnight in an incubator. The media was removed from each well, replaced with 1 mg/L each (GT)_n_-SWCNT diluted in media, and incubated for 30 minutes (pulsed) to allow for internalization into the cells. After this pulse, the SWCNT-containing media was removed, the cells were rinsed 3X with sterile PBS (Gibco) and fresh media was replenished. Well plates were mounted on the hyperspectral microscope to obtain broadband images, transmitted light images, and fluorescence hyperspectral images at each given time point. Hyperspectral data were processed and extracted using custom codes written with Matlab software. All Gaussian curve fits were generated using OriginPro 2018.

### Confocal Raman Microscopy

Cells were seeded into 35mm glass bottom microwell dishes (MatTek) to a final concentration of 500,000 cells/ dish and allowed to culture overnight in incubator. The media was removed from each well, replaced with 1 mg/L (GT)_6_-SWCNT or (GT)_30_-SWCNT diluted in media, and pulsed for 30 minutes to allow internalization into the cells. The SWCNT-containing media was removed, the cells were rinsed 3X with sterile PBS (Gibco), and fresh media was replenished. The 0-hour samples were immediately fixed using 4% paraformaldehyde in PBS for 10 minutes, rinsed 3X with PBS, and covered with PBS to retain an aqueous environment during imaging. The 24-hour samples were later fixed using the same procedure. The cells were imaged using an inverted WiTec Alpha300 R confocal Raman microscope (WiTec, Germany) equipped with a Nikon CFI-Achro 60x/0.8 air objective, a 785nm laser source set to 35mW sample power, and collected with a CCD detector through a 600 lines/mm grating. The Raman spectra were obtained in 0.5×0.5 μm intervals with 1s integration time to construct hyperspectral Raman area scans of cellular regions. A calibration curve was obtained by recording spectra of known SWCNT concentrations serially diluted in a single pixel volume with identical acquisition settings. Each spectrum was averaged over 20 scans. A global background subtraction and cosmic-ray removal was performed using Witec Control 5.0 software on all acquired confocal Raman data and G-band maximum intensities were extracted and correlated with known concentrations to produce a linear curve fit using OriginPro 2018 analysis software. The cellular SWCNT concentration data were produced by relating the G-band linear equations to each SWCNT-containing pixel and intracellular concentrations were obtained in individual cell ROIs with the correlation 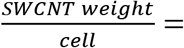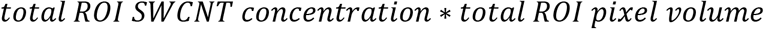.

### Solution-Based Fluorescence Dequenching Assay

Cy3-(GT)_6_- or Cy3-(GT)_30_ oligonucleotides were purchased from Integrated DNA Technologies and used in the creation of DNA-SWCNTs (see above). After ultrasonication and ultracentrifugation, the Cy3-DNA-SWCNTs were filtered 3 times using 100 kDa Amicon centrifuge filters (Millipore) to remove free Cy3-DNA from solution, diluted to 2.5 mg/L, and 1 mL was placed in a plastic cuvette under magnetic stirring. The fluorescence intensity of each sample was obtained in 1-second intervals for 3 minutes using a Perkin Elmer LS 55 fluorescence spectrometer set to 532 nm excitation and 569 nm emission with 3 nm bandwidth. A 10 μL aliquot of a 10% sodium deoxycholate solution (Sigma-Aldrich) was spiked into the Cy3-DNA-SWCNTs after a baseline intensity was established for a final concentration of 0.1% SDC in order to temporally displace the Cy3-DNA from the SWCNTs as previously described [28].

### Visible Fluorescence Microscopy in Live Cells

Cy3-(GT)_6_-SWCNTs and Cy3-(GT)_30_-SWCNTs were first filtered 3 times using 100kDa Amicon centrifuge filters (Millipore) to remove free Cy3-DNA from solution. The cells were seeded onto 35 mm glass-bottom petri dishes (MatTek) to a final concentration of 500,000 cells/dish and allowed to culture overnight in an incubator. The media was removed from each well, replaced with 1 mg/L of filtered Cy3-(GT)_6_-SWCNTs or Cy3-(GT)_30_-SWCNTs diluted in media, and incubated for 30 minutes to allow internalization into the cells. The SWCNT-containing media was removed, the cells were rinsed 3X with sterile PBS (Gibco), and fresh media was replenished for each sample. The petri dishes were mounted in a stage incubator (Okolab) on an Olympus IX-73 inverted microscope with a UApo N 100x/1.49 oil immersion objective for epifluorescence imaging with a U-HGLGPS excitation source (Olympus) filtered through a Cy3 filter cube. The fluorescence images were analyzed by extracting average fluorescence intensity values of individual cell ROIs using ImageJ.

### Statistical Analysis

All statistical measures for hypothesis testing were carried out using two-sample two-tailed unequal variance t-tests in Microsoft Office Excel 2016. All curve fitting and related statistics were performed in OriginPro 2018.

## Acknowledgements

This work was supported by the RI-INBRE Early Career Development Award Grant #P20GM103430 from the National Institute of General Medical Sciences of the National Institutes of Health, the Rhode Island Foundation – Medical Research Fund, and the URI College of Engineering. The confocal Raman data was acquired at the RI Consortium for Nanoscience and Nanotechnology, a URI College of Engineering core facility partially funded by the National Science Foundation EPSCoR, Cooperative Agreement #OIA-1655221.

