## Supporting Information for "Oligonucleotide Length Determines Intracellular Stability of DNA-Wrapped Carbon Nanotubes"

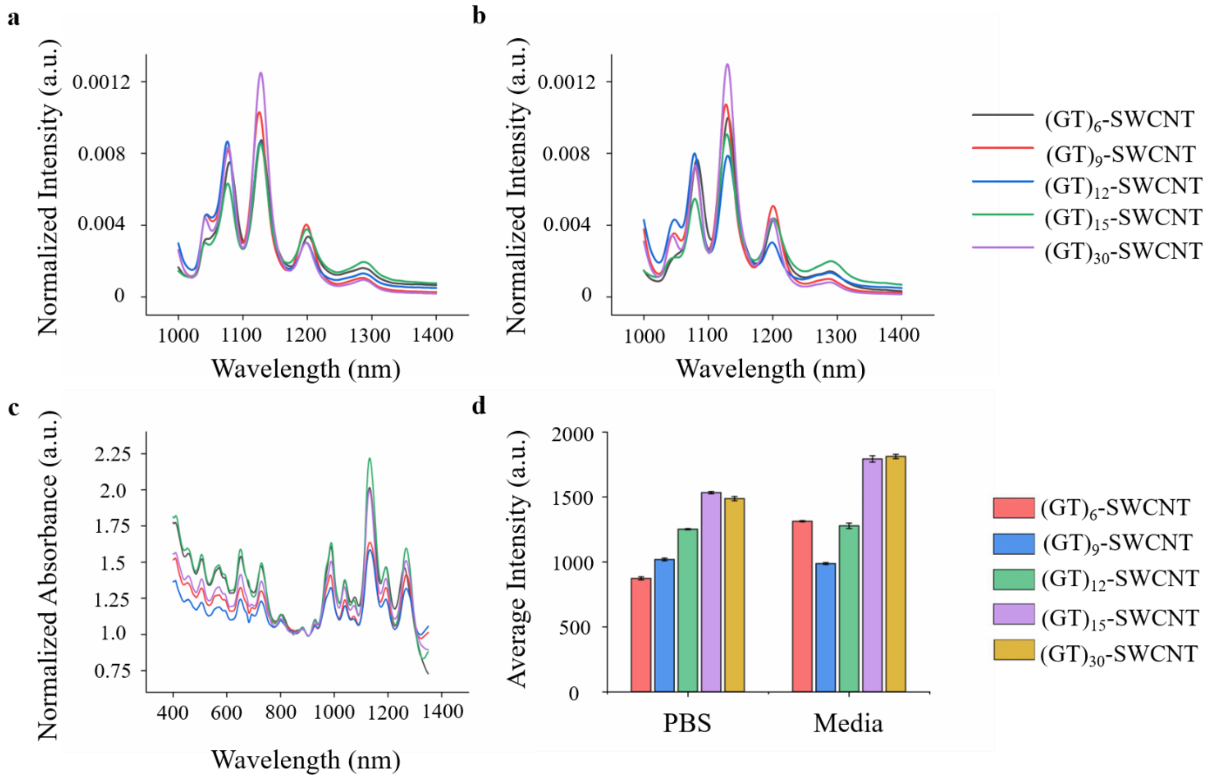


**Figure S1.** Solution-based optical characterizations of DNA-SWCNTs. Fluorescence spectra for all $(GT)_n$-SWCNTs diluted to 1 mg/L in **(a)** PBS or **(b)** cell culture media normalized to area under the curve. **(c)** Absorbance spectra for all $(GT)_n$-SWCNTs normalized to the absorbance at 910 nm. **(d)** Integrated fluorescence intensity of all $(GT)_n$-SWCNTs diluted to 1 mg/L in PBS or media represented as average ± s.d. n = 4.

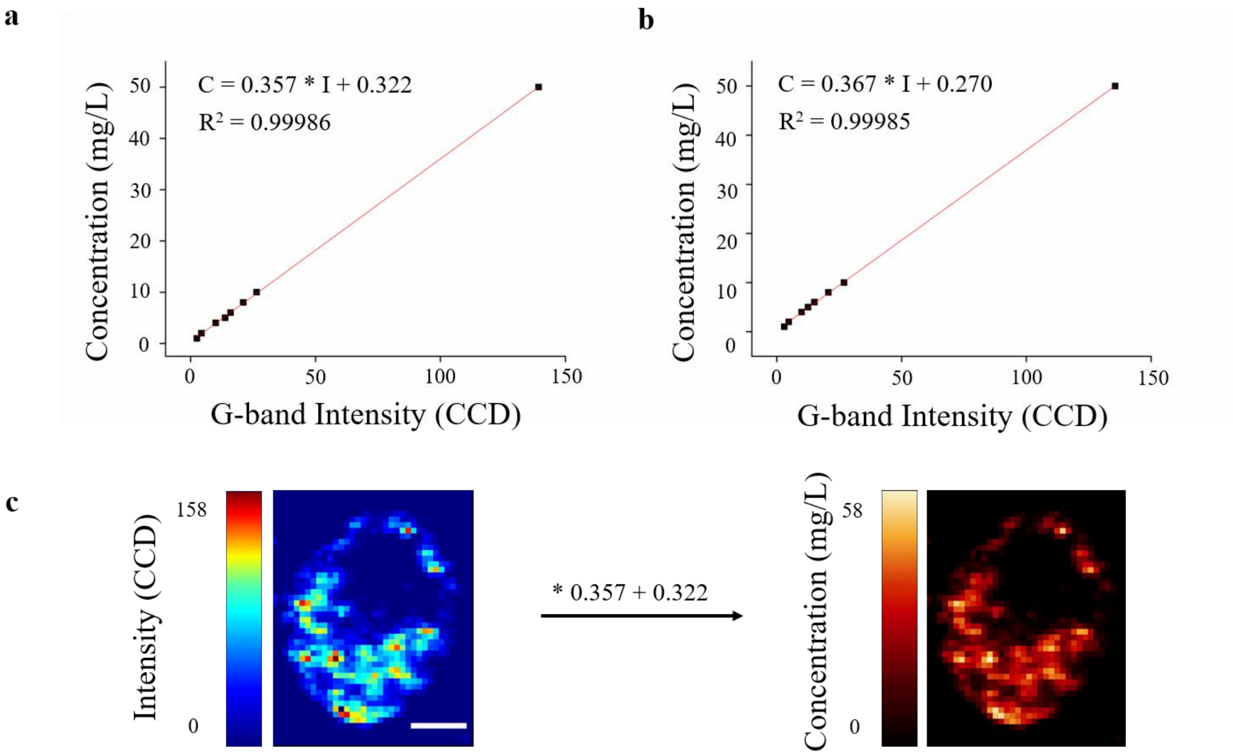


**Figure S2.** Confocal Raman concentration-intensity calibration. Linear fits of G-band intensity versus known **(a)** $(GT)_6$-SWCNT and **(b)** $(GT)_{30}$-SWCNT concentrations. **(c)** Example of cell ROI CCD intensity scale converted to concentration using linear fit equation.

| Peak Emission Energy Shifts | | | | | | | |
|---|---|---|---|---|---|---|---|
| Chirality ($n$,$m$) | ssDNA | 0h | | 6h | | 24h | |
| | | Average (meV) | ± s.e. | Average (meV) | ± s.e. | Average (meV) | ± s.e. |
| (10,2) | $(GT)_6$ | -1.518 | 0.392 | 3.338 | 1.046 | 5.084 | 0.307 |
| | $(GT)_9$ | -0.278 | 0.053 | 0.595 | 0.510 | -2.275 | 0.873 |
| | $(GT)_{12}$ | -1.046 | 0.379 | -2.761 | 0.155 | -3.237 | 0.267 |
| | $(GT)_{15}$ | -0.268 | 0.341 | -1.500 | 0.611 | -2.166 | 0.441 |
| | $(GT)_{30}$ | -0.427 | 0.063 | -1.427 | 0.252 | -2.452 | 0.164 |
| (9,4) | $(GT)_6$ | -2.065 | 0.183 | 5.706 | 0.253 | 0.694 | 0.047 |
| | $(GT)_9$ | -3.130 | 0.313 | -2.105 | 0.172 | -5.008 | 0.233 |
| | $(GT)_{12}$ | -6.382 | 0.627 | -3.088 | 0.337 | -3.239 | 0.164 |
| | $(GT)_{15}$ | -3.354 | 0.058 | -3.856 | 0.218 | -4.854 | 0.065 |
| | $(GT)_{30}$ | -3.099 | 0.208 | -4.313 | 0.346 | -4.768 | 0.084 |
| (8,6) | $(GT)_6$ | -1.312 | 0.109 | 4.927 | 0.463 | -0.019 | 0.230 |
| | $(GT)_9$ | -1.756 | 0.128 | -2.414 | 0.249 | -3.538 | 0.052 |
| | $(GT)_{12}$ | -2.528 | 0.347 | -4.313 | 0.412 | -5.063 | 0.332 |
| | $(GT)_{15}$ | -1.909 | 0.144 | -3.185 | 0.225 | -3.885 | 0.156 |
| | $(GT)_{30}$ | -1.437 | 0.287 | -2.851 | 0.273 | -3.409 | 0.248 |
| (8,7) | $(GT)_6$ | -4.360 | 1.043 | -3.435 | 1.948 | -3.784 | 1.341 |
| | $(GT)_9$ | -1.960 | 0.582 | -6.161 | 2.107 | -3.434 | 1.705 |
| | $(GT)_{12}$ | -3.085 | 1.301 | -1.402 | 1.464 | -2.688 | 0.983 |
| | $(GT)_{15}$ | -3.339 | 0.714 | -3.756 | 0.683 | -4.209 | 0.859 |
| | $(GT)_{30}$ | -4.699 | 0.642 | -7.720 | 0.920 | -5.544 | 0.441 |

**Table S1.** Table of calculated intracellular peak emission energy shifts for all examined $(GT)_n$-SWCNTs. Gaussian functions were fitted to each spectrum to obtain peak emission energies. Emission peaks in cell culture media were subtracted from each intracellular emission energy to determine shift values. n = 3.
